# A matter of food and substrain: obesogenic diets induce differential severity of cardiac remodeling and diastolic dysfunction in C57Bl/6J and C57Bl/6N substrains

**DOI:** 10.1101/2024.03.15.584713

**Authors:** Lorena Cascarano, Hrag Esfahani, Pierre Michel, Caroline Bouzin, Chantal Dessy, Jean-Luc Balligand, Lauriane Y.M. Michel

**Affiliations:** Institute of Experimental and Clinical Research (IREC), Pharmacology and Therapeutics (FATH), Cliniques Universitaires St Luc and Université catholique de Louvain (UCLouvain), 1200 Brussels, Belgium; 2IP-IREC Imaging Platform, Institute of Experimental and Clinical Research (IREC), Université catholique de Louvain (UCLouvain), RRID:SCR_023378, 1200 Brussels, Belgium

## Abstract

The prevalence of metabolic syndrome in cardiac diseases such as heart failure with preserved ejection fraction (HFpEF) prompts the scientific community to investigate its adverse effects on cardiac function and remodeling and its associated mechanisms. However, the choice of a preclinical model of obesity-induced cardiac remodeling has proven more challenging with inconsistencies often found in very similar mouse models. Here, we investigated the implication of both genetic background of mouse substrains as well as diet composition to identify a suitable model of diet-induced cardiac alterations. C57Bl/6J and C57Bl/6N male mice were subjected to distinct obesogenic diets consisting of high-fat and moderate-sucrose content (HF-S) or High-Sucrose and moderate lipid content (F-HS) versus matching control diets. 5-month dietary intervention with obesogenic diets induced weight gain, adipocyte hypertrophy and increased visceral and subcutaneous fat mass in both substrains. Obese mice showed similar impairment of glucose disposition and insulin tolerance among genotypes, both strains developing insulin resistance within two months. However, echocardiographic follow-up and histological analysis confirmed that HF-S diet increases cardiac hypertrophy, interstitial fibrosis as well as le atrial area in the C57Bl/6J strain only. On the contrary C57Bl/6N exhibit cardiac eccentric remodeling under control diets, possibly owing to a genetic mutation in the myosin light chain kinase 3 (*Mylk3)* gene, specific to this substrain, which was not further enhanced under obesogenic diets. Altogether, the present results highlight the importance of carefully selecting the suitable mouse strain and diets to model diet-induced cardiac remodeling. In this regard, C57Bl/6J mice develop significant cardiac remodeling in response to HF-S, and seem a suitable model for cardiometabolic disease.

## INTRODUCTION

Metabolic Syndrome affects over a billion persons worldwide^1^ with a prevalence expected to steadily increase in the coming decades^2^. Metabolic syndrome is primarily caused by obesity and associated metabolic dysfunction which promote the development of cardiovascular diseases^3^. Major interrelated risk factors include abdominal (visceral) obesity, impaired glucose tolerance, hypertriglyceridemia, hypertension and decreased HDL. Since this description, cardiovascular disease prevention has focused primarily around atherogenic and hypertensive risks^4^ and around nutrient overload and diabetic glucose imbalance causing diabetic cardiomyopathy^5,6^. Nevertheless, the strikingly high prevalence of metabolic syndrome in cardiac pathologies such as in heart failure with preserved ejection fraction (HFpEF) (ranging from 25 to 60%^7-9^) and the adverse clinical phenotype^10^ urge the scientific community to investigate the early steps of metabolic syndrome, including the incipient obesity and cardiac remodeling. Importantly, this process appears long before atherogenic complications, coronary perfusion damage or diabetes-induced cardiomyopathy, suggesting that early metabolic alterations and ensuing oxidative stress and inflammatory state may initiate cardiac remodeling^9,11^.

Mechanisms underlying adverse cardiac alterations have not yet been thoroughly investigated, owing to the lack of a reliable pre-clinical animal model of diet-induced cardiac remodeling in particular in the mouse, a model widely used to test genetic modifications. Indeed, the development of a mouse model of nutrient overload and obesity-induced cardiac remodeling has proven challenging with inconsistencies often found in very similar mouse models. In this regard, food composition as well as variations among inbred strains regarding sensitivity to nutrient overload have been repeatedly overlooked in experimental settings, precluding consistent results under the commonly called “high-fat diet” (HFD) regimen. Across the classic laboratory inbred strains, an impressively large number of genomic variations has been catalogued^12^. In parallel, previous studies have observed that while most mouse strains develop metabolic defects when subjected to a obesogenic diet, the genetic background of the different strains leads to inherent differences in metabolic parameters^13,14^. Later studies confirmed that this also holds true among lines deriving from a similar strain, also known as substrains^15^.

Among the commonly used mouse strains, the inbred C57Bl/6 strain prevail as the most widely used worldwide. Derived from the C57BL line established in 1921 by Dr. C.C. Little the C57Bl/6 line was introduced at the Jackson Laboratory in 1948 and was passed to the National Institutes of Health (NIH) in 1951^16,17^. Since this date, inbred colonies of C57Bl/6 mice have been separately maintained in both laboratories giving rise to C57Bl/6J and C57Bl/6N substrains^18^. Genome sequence comparisons of C57Bl/6J and C57Bl/6N revealed 34 SNPs and 2 indels as well as 15 structural variants between the two substrains^19^ leading to a wide variety of phenotypic differences in the response of each substrain to HFD.

These differences have been previously examined with respect to metabolic responses with C57Bl/6J clearing glucose less rapidly than other strains^20^. Later studies attributed this effect to lower insulin secretion in C57Bl/6J due to reduced responsiveness of β-cells to glucose^15,21,22^.Regarding adiposity C57Bl/6J strain was originally reported to gain more weight under HFD, and thus has been considered more susceptible to diet-induced obesity^23^. Therefore, this substrain has been widely used in metabolic research, although the difference in weight gain is not consistently supported in the literature^24^. Many addional differences have been idenfied between susbtrains in non-metabolic parameters such as responses to alcohol^25^, susceptibility to chronic pancreatitis^26^, systolic arterial pressure and pulse rate^19^. However, no study so far examined the cardiac phenotype among substrains under obesogenic diets and the possible disparities when facing nutrient overload.

The validation of an adequate substrain to model diet-induced cardiac remodeling and possibly early steps of diastolic dysfunction would represent a considerable advance to further dissect the implication of diet in the early pathogenesis of current cardiac pathologies such as HFpEF and to investigate paracrine and endocrine communication between organs under obesogenic diets. Therefore, in the present study, we evaluate the sensitivity to obesity-induced cardiac remodeling of the C57Bl/6J and C57Bl/6N substrains (referred as 6J and 6N, respectively) by a side-by-side comparison of metabolic parameters, adipose tissue remodeling and an extensive characterization of cardiac remodeling and cardiac function. To this aim, both substrains were subjected to two distinct types of obesogenic diets and their matching control diets during 5 months.

## MATERIALS AND METHODS

### Mice

32 C57Bl/6JRj and 32 C57Bl/6NRj male mice (9-week-old) were obtained from Janvier Labs (Le Genest Saint-Isle, France). Animals were housed in 12h/12h day/night cycle and had a free access to water and chow diet (except when it is specified). Experiments were restricted to male to avoid confounding effects of hormonal status in female. Both experiments (Bl6J and Bl6N) were realized separately but during the year period (between October and December). 9-week mice were randomly and equally divided into four groups. After 3 weeks of acclimation, each group was assigned a specific diet; chow diet (SAFE diets, Augy, France), control diet (Research diets, New Brunswick, USA), high-fat - sucrose (HF-S) diet (SAFE diets, Augy, France), Fat - High-Sucrose (F-HS) diet (Research diets, New Brunswick, USA). The compositions of the diets are detailed in Table 1. Mice were fed during 22 weeks with these diets before being sacrificed to collect adipose tissues and organs. Body weight was measured prior to the start of the diets and was monitored each week during the first month and every other week until the end of the diet period. This study was carried out in accordance with the Guide for the Care and Use of Laboratory Animals published by the U.S. National Institutes of Health (NIH) and the European Directive 2010/63/EU and were approved by local ethical committees.

**Table 1.** Nutritional composition of the diets.

|  | SAFE |  | Research diets |  |
| --- | --- | --- | --- | --- |
|  | Chow diet | HF-S diet | Control diet | F-HS diet |
| Reference | A04 | U8978<br>version 19 | D14042701 | D12079B |
|  | Energy content (% KCal) |  |  |  |
| Fat | 8,4 | 58,8 | 10 | 40 |
| Carbohydrate | 72,2 | 26,8 | 73 | 43 |
| of which sucrose | 0 | 13 | 0 | 29 |
| Protein | 19,3 | 14,3 | 17 | 17 |
| Fibers (%gm) | 4% crude | <0,5% crude | 4% Cellulose | 5% Cellulose |
|  | Caloric content (Kcal/kg) |  |  |  |
|  | 3339 | 5497 | 3915 | 4679 |

### Echocardiography

Mice were anesthetized by 1-3% isoflurane inhalation and were submitted to a two-dimensional echocardiography with a Vevo 2100 Imaging system (VisualSonics, Toronto, ON, Canada). LV parasternal long axis view was recorded in B-mode to measure LV internal volumes at end of diastole (LVEDv) and systole (LVESv), used to calculate the ejection fraction (EF; Bmode). M-mode view was performed to measure LV posterior wall (PW) and interventricular septum (IVS) thicknesses, LV internal dimensions (LVID), allowing the assessment of the fractional shortening (FS). LV mass was calculated as 1,053*[(LVIDd + PWd + IVSd)^3^ - LVIDd^3^]. Measurements were carried out by the same operator. Echocardiographic tracings were performed in a blinded fashion using Vevo LAB software.

### Glucose and insulin tolerance tests

Intraperitoneal glucose and insulin tolerance tests (IP-GTT and IP-ITT) were performed at baseline and every month during the diet period. IPGTT was assessed after 6h-fasting by measuring tail blood glucose at 0, 15, 30, 60 and 120min in response to a 2g/kg glucose injection using test strips with the associated Contour Next glucose meters (Ascensia). The same method was applied for the IP-ITT with a fasting time of 4h before administration of 0,5U/kg of insulin. Data are expressed as plasma glucose (mg/dl) for IP-GTT and as % of basal glycemia for IP-ITT.

### Tissue collection and dissection

During the sacrifice, fat pads were collected from inguinal subcutaneous adipose tissue, epididymal adipose tissue and pericardial adipose tissue; tissues were harvested in a consistent manner (inguinal and epididymal right fat pad). Heart was stopped in diastole by immersion in KCl 50mM, was then weighed and right ventricle excised to measure left ventricular (LV) weight. Lungs were collected and directly weighted (wet weight) and then dried at 60°C °C for 24hours to measure dry weight. LV as well as pieces of the subcutaneous inguinal and visceral epididymal adipose tissues were embedded in paraffin for histological analysis.

### Adipocyte isolation

Around 300mg of inguinal adipose tissue were collected during sacrifice, cut in small pieces and digested at 37°C with collagenase A at 1mg/kg (Sigma-Roche, #10103578001) in Krebs - 1% BSA buffer during roughly 20 minutes. After 3 washing steps with Krebs - 1% BSA (5min centrifugation at 1500rpm), adipocytes corresponding to the floating cell layer were collected and lysed with TriReagent (MRC Inc, OH, US) prior to RNA extraction.

### RNA extraction and real-time reverse transcription–polymerase chain reaction

Total RNA was extracted from murine tissues using TRI-reagent (Fermentas, Alost, BE) and subsequent homogenization with Precellys Evolution (Bertin technologies) and extraction according to standard protocols. Genomic DNA contamination was prevented by treatment with RQ1 RNase-free DNase (Promega, Inc.). Subsequently, 1μg of each sample RNA was reverse-transcribed with RevertAid Reverse Transcriptase using OligodT and random hexamers (Thermo Scientific).

Transcript expression was assessed using low ROX Takyon (Eurogentec) on 1:25 diluted cDNA with the CFX96 real-time system (BIORAD, Laboratories GmbH, Vienna, Austria). Results are expressed as 2^-ΔΔCt^ (Livak method) normalized on housekeeping gene expression HPRT (hypoxanthine phosphoribosyl transferase). The exon-overlapping amplicons were amplified using the following calibrated primers sets: Adiponectin forward 5’-GGC-TCT-GTG-CTG-CTC-CAT-CT-3’; CIDEA 5’-CAT-ACA-TGC-TCC-GAG-TAC-TGG-3’; TBX1 5’-TGG-GAC-GAG-TTC-AAT-CAG-C-3’; TNFα 5’-CCC-TCA-CAC-TCA-GAT-CAT-CTT-CT-3’; NOS2 5’-GGA-GTG-ACG-GCA-AAC-ATG-ACT-3’; IL6 5’--CTG-CAA-GAG-ACT-TCC-ATC-CAG-3’.

### Histology and morphometry

Left ventricle as well as inguinal and epididymal adipose tissue sections were fixed overnight in 4% formaldehyde. The paraffin-embedded tissues were cut into 5μm-thick slices. After deparaffination and rehydration, tissue sections were stained with rhodamine-conjugated–wheat germ agglutinin (WGA; Vector RL-1022; vector labs, 2h-incubation, 1/300), Perilipin-1 (abcam; ab3526), F4/80 (Cell signaling; 70076), CD86 (Cell signaling; 19589) or with Sirius Red (SR, 2h-incubation) at room temperature using standard techniques. Slides were mounted with the Sakura (SR) or manually (other stainings) and pictures were acquired with a Panoramic ScanII (SR) and AxioScan 21 (other stainings) and analyzed in a blinded fashion using Visiopharm software.

### Statistical analysis

All results are expressed as means ± SEM and were then analyzed by unpaired Student’s t test, one-way analysis of variance (ANOVA), two-way ANOVA, or two-way ANOVA for repeated measures where appropriate. In the absence of a normal distribution, nonparametric Krukal-Wallis or Mann-Whitney test were used. P < 0.05 was considered statistically significant. Statistical analyses and data visualization were realized using GraphPad Prism (GraphPad Software, San Diego, CA).

## RESULTS

To compare the cardio-metabolic phenotype between C57Bl/6J and C57Bl/6N substrains (referred thereafter as 6J and 6N, respectively), 32 mice from each genotype (12-week old) were fed either a HF-S (High-Fat – Sucrose ; 59% Kcal from lipids, 13% Kcal from sucrose))(n=8) compared with its respective littermates under regular chow diet (CD; n=8)(composition in **table 1**) or F-HS (Diet containing fat and high sucrose; 40% Kcal from lipids, 29% Kcal from sucrose) (n=8) diet compared with animals under control diet with matching refined fibers (CTL diet; n=8) (**table 1**) over 22 weeks (**Fig. 1**). All mice underwent cardiac echocardiography at the start and at 4,8,12, 16, 20 weeks of the diet and were sacrificed at 22 weeks. Glucose and insulin tolerance tests and weight measurements were performed at the same time intervals. After sacrifice, visceral (epididymal) and subcutaneous (inguinal) adipose tissues were collected and either fixed for immunostaining, frozen for RNA extraction, or dissociated for isolation of differentiated adipocytes. Hearts and lungs were collected, weighted and dissociated or fixed for similar analyses.

**Figure 1.**
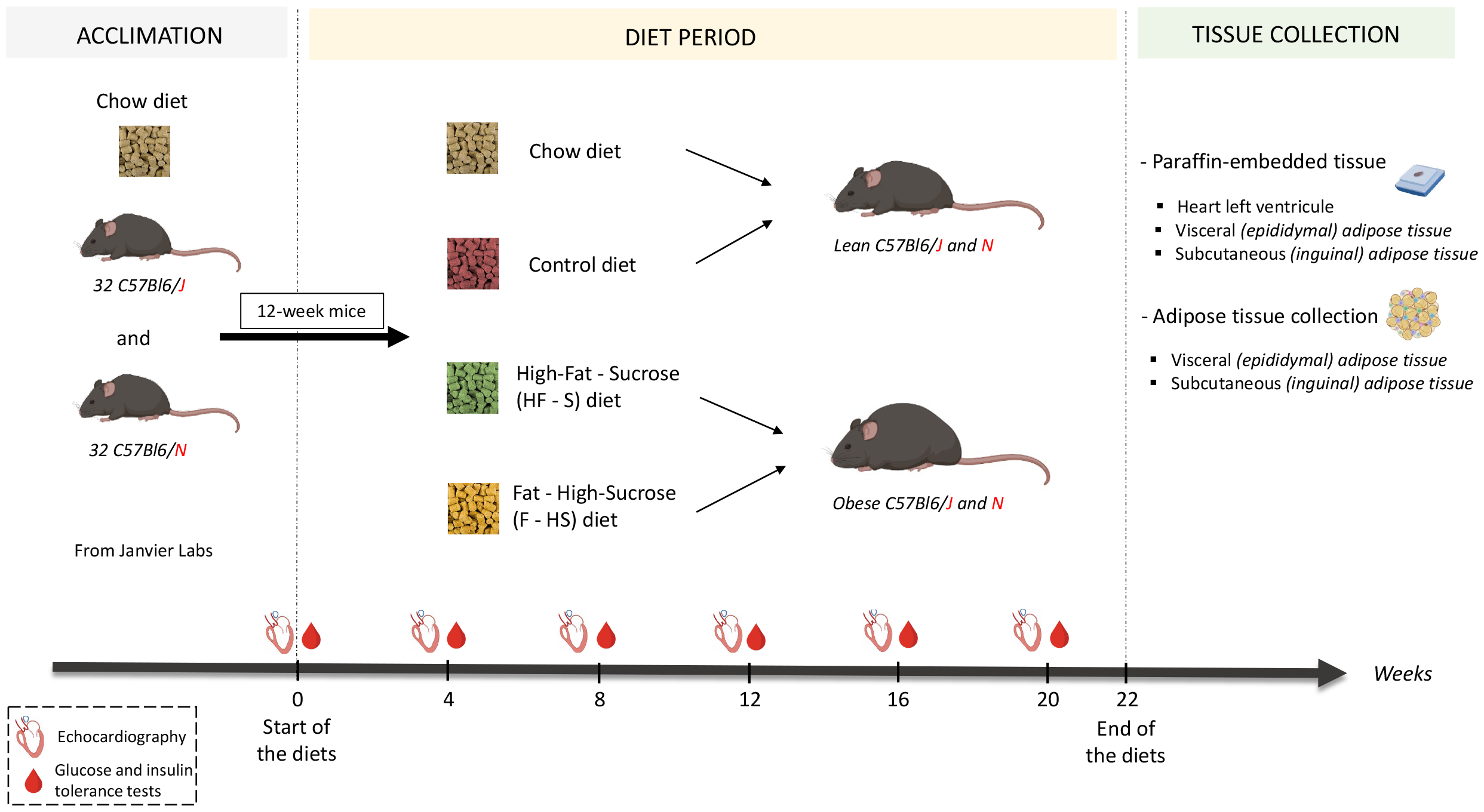
Experimental setup.

### Comparative changes in adipose tissue and metabolic parameters under obesogenic diets among substrains

Both HF-S and F-HS diets induced a progressive weight gain in both strains compared with control diets. Both strains gained more weight under HF-S than F-HS (**Fig. 2A&2I**). Notably, at the same 12-week age, baseline weight was higher in 6N, which also gained weight more rapidly under HF-S than 6J until maximum weight at 22 weeks which was comparable between the 2 strains. This was reflected by similar increases in inguinal and epididymal fat mass and in adipocyte size in the 2 strains (**Fig. 2B-2E & 2J-2M**). In isolated adipocytes, mRNA expression of adipose marker adiponectin as well as markers of beige thermogenic adipocytes CIDEA and TBX1 were similarly reduced in both strains under HF-S and F-HS diets (**Fig. 2F-2H & 2N-2P**).

**Figure 2.**
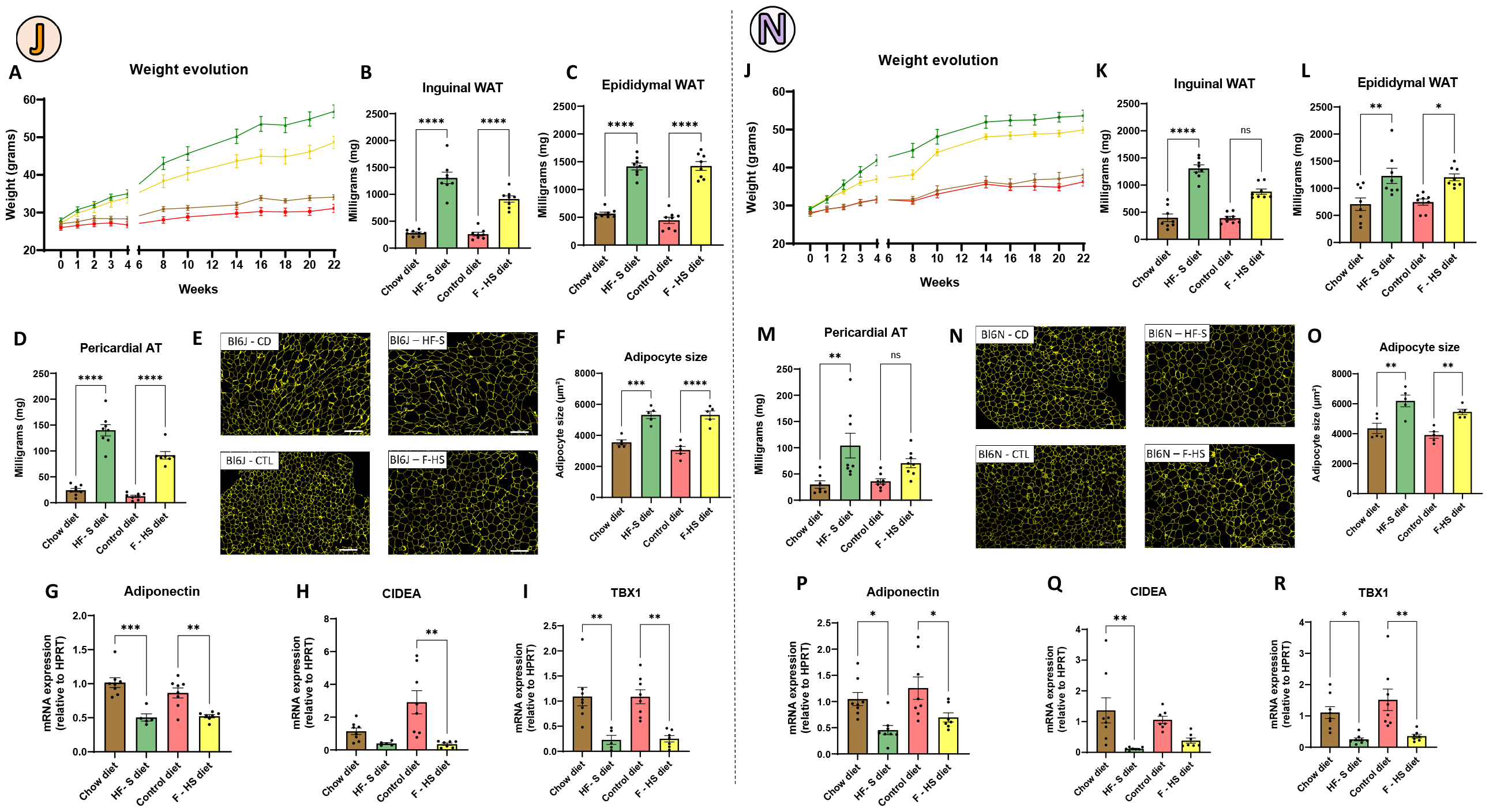
Obesogenic diets induce comparable changes in weight and adipose tissue characteristics. Weight evolution of the C57Bl/6J (A) and C57Bl/6N (J) mice treated with chow diet (brown), control diet (red), HF – S (green) and F – HS (yellow)(n = 8 mice per group). Weight of Inguinal, epididymal and pericardial adipose tissue in C57Bl/6J (B-D) and C57Bl/6N (K-M). Adipocyte area determined by perilipin-1 immunostaining in C57Bl/6J (E-F) and C57Bl/6N (N-O)(scale bar 200μm). Transcriptional determination of Adiponectin, Cidea and Tbx1 expression in C57Bl/6J (G-I) and C57Bl/6N (P-R). Data indicate the mean +/-SEM and were analyzed by a 1-way ANOVA.

Diet-induced changes in glucose and insulin tolerance were measured in the 2 strains over time (M0 to M5). Under both control diets, the maximal glucose concentration and AUC for glucose after i.p. glucose load were higher in 6J than 6N mice (1047,0 +/-49,1 *vs* 775,8 +/-64,1, *p* < 0,0001)(**Suppl. Fig. 1A&1E**). Both strains responded with higher maximal glucose and AUC after glucose load after two months of both HF-S and F-HS diets, with HF-S producing more marked increases (**Suppl. Fig. 1B-F**). Notably, at M5, the difference between the curves for HF-S diet and control diets tended to attenuate, probably due to a combined effect of both degradation of glycemic tolerance with aging under control diets and increased production of insulin with progressive obesity under HF diet. This metabolic degradation was confirmed by measurements of insulin tolerance over time (**Fig. 3**). At baseline, the maximal decrease in glycemia after injection of insulin was more marked in 6J vs 6N mice (362,0 +/-21,8 vs 408,4 +/-20,4, p<0,0001); starting at M2, this difference was more pronounced between 6J and 6N under control diet; however, in both strains, the effect of insulin was blunted under HF-S and F-HS diets, with an almost abolished response to insulin under HF-S in both strains.

**Figure 3.**
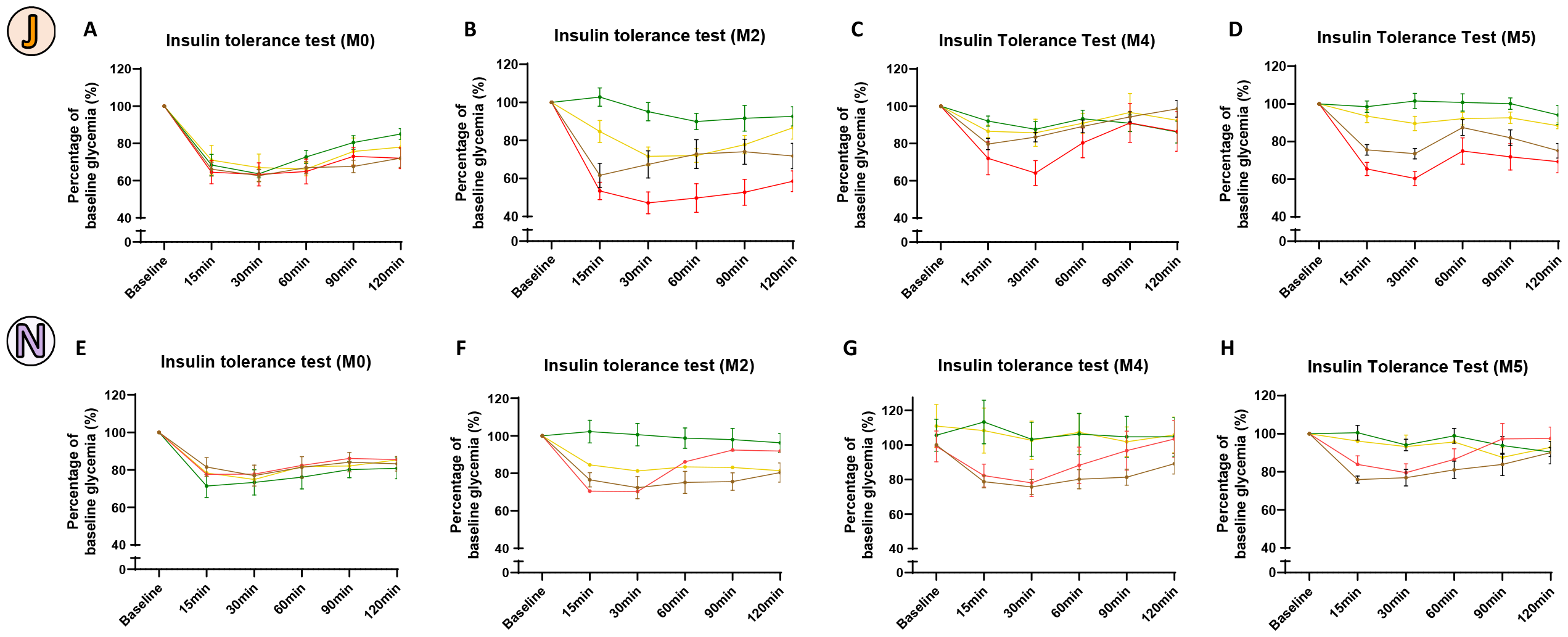
Substrain sensitivity to insulin under obesogenic diets. Intraperitoneal insulin tolerance test (0,5U/kg) of the C57Bl/6J mice (A-D) and C57Bl/6N mice (E-H) treated with chow diet (brown), control diet (red), HF – S (green) and F – HS (yellow) at baseline (A-E) and after 2 (B-F), 4 (C-G) and 5 (D-H) months of treatment with the diets. Data indicate the mean +/-SEM (n = 8 mice per group) and were analysed by a 2-way ANOVA for repeated-measures.

### Differential severity in cardiac remodeling depends on substrains and diets

Next, the effect of diets on the cardiac phenotype was compared between 6J and 6N mice. **Fig. 4** and **Fig. 5** and **Suppl. Tables 1 and 2** show the changes in heart and left ventricular (LV) remodeling and function over time under control or obesogenic diets. Heart rate and ejection fraction were comparable between control diets and obesogenic diets under both strains at M5; however, 6N mice exhibited higher indexes of total heart (9,3 +/-0,4 mg vs 7,8 +/-0,2 mg) and LV mass (5,9 +/-0,4 mg vs 4,7 +/-0,2 mg), as well as left atrial dimensions (3,50 +/-0,20 mm^2^ vs 2,17 +/-0,05 mm^2^) compared with 6J under chow diet (**Fig. 5C-E**). Notably, echocardiographic measurements at baseline (before obesogenic diets) showed enlarged LV dimensions in 6N compared to 6J mice (**Suppl. Fig. 2E-F**). Only 6J mice developed a significant LV hypertrophy (increased LV mass and relative wall thickness by echocardiography) specifically under HF-S diet (**Fig. 4C-E**), whereas 6N mice did not further increase these parameters beyond levels observed under control diets. Increases in cardiac and LV hypertrophy (total heart mass and LV mass normalized to tibial length) under HF-S diet were confirmed by gravidometric analysis after sacrifice (M5) in 6J only (**Fig. 4F-G**). Histo-morphometric analysis in fixed LV sections showed increased transverse size of cardiac myocytes (WGA staining)(**Fig. 6C-D**) as well as increased interstitial fibrosis (Sirius red staining) (**Fig. 6A-B**)in hearts from 6J mice under HF-S diet at M5. This LV remodeling was accompanied with increased left atrial size (**Fig. 4E**) by echocardiography as well as increased lung weight (**Suppl. Fig. 2A-D**), suggestive of increased LV filling pressures, at M5 under HF-S diet compared with control diet in 6J, but not 6N, in which these parameters were already elevated under control diet, suggesting some other constitutive element driving cardiac remodeling in 6N mice (**Fig. 5E**). Such remodeling was not accompanied with interstitial fibrosis, even under HF-S diet, contrary to 6J mice (**Fig. 6B**), however an increase in LV internal dimensions in systole and diastole was observed in 6N compared to 6J (**Suppl. Fig. 2E-F**).

**Figure 4.**
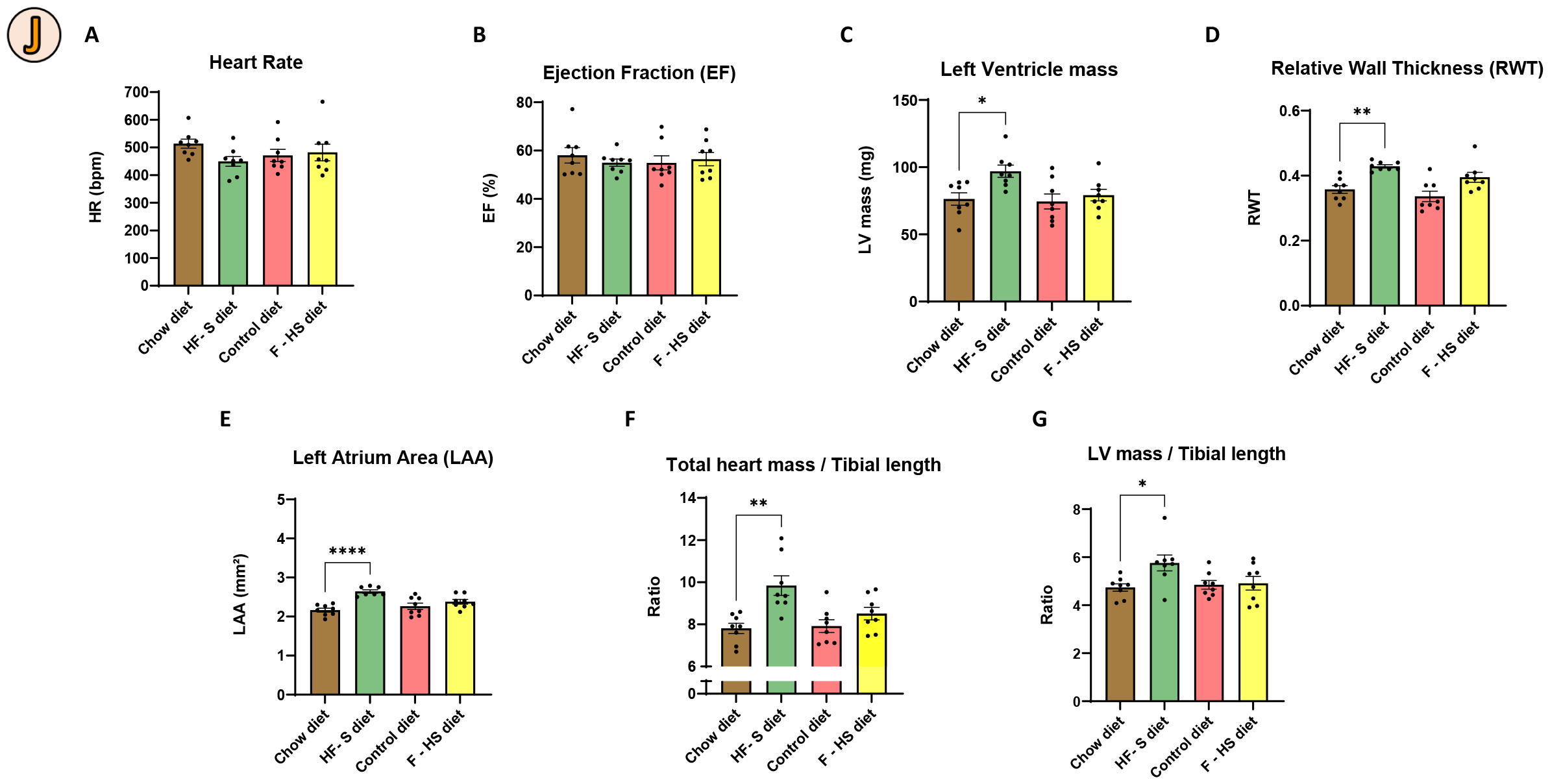
Effect of the diets on the cardiac and morphometric parameters of C57Bl/6J mice. Echocardiographic analysis of heart rate (A), ejection fraction (B), le ventricule mass (C), relative wall thickness (D), le atrium area (E) and ratio of the total heart (F) and le ventricule (LV) mass (G) on the tibial length measured after the sacrifice of the C57Bl/6J mice treated with chow diet (brown), control diet (red), HF – S (green) and F – HS (yellow) after 22 weeks of treatments. Data indicate the mean +/-SEM (n = 8 mice per group) and were analysed by a one-way ANOVA test.

**Figure 5.**
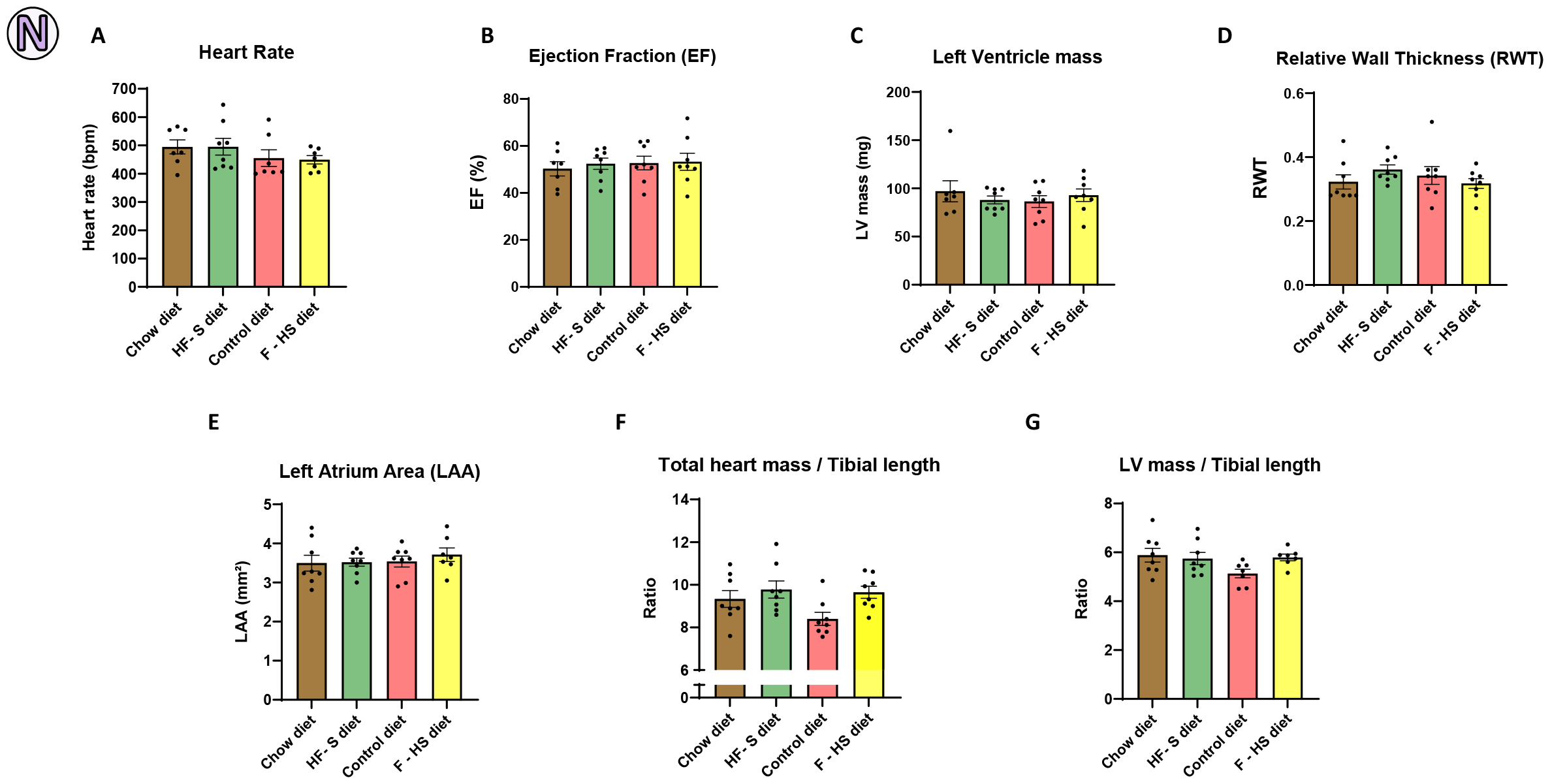
Effect of the diets on the cardiac and morphometric parameters of C57Bl/6N mice. Echocardiographic analysis of heart rate (A), ejection fraction (B), left ventricule mass (C), relative wall thickness (D), le atrium area (E) and ratio of the total heart (F) and le ventricule (LV) mass (G) on the tibial length measured after the sacrifice of the C57Bl/6N mice treated with chow diet (brown), control diet (red), HF – S (green) and F – HS (yellow) after 22 weeks of treatments. Data indicate the mean +/-SEM (n = 8 mice per group) and were analysed by a one-way ANOVA test.

**Figure 6.**
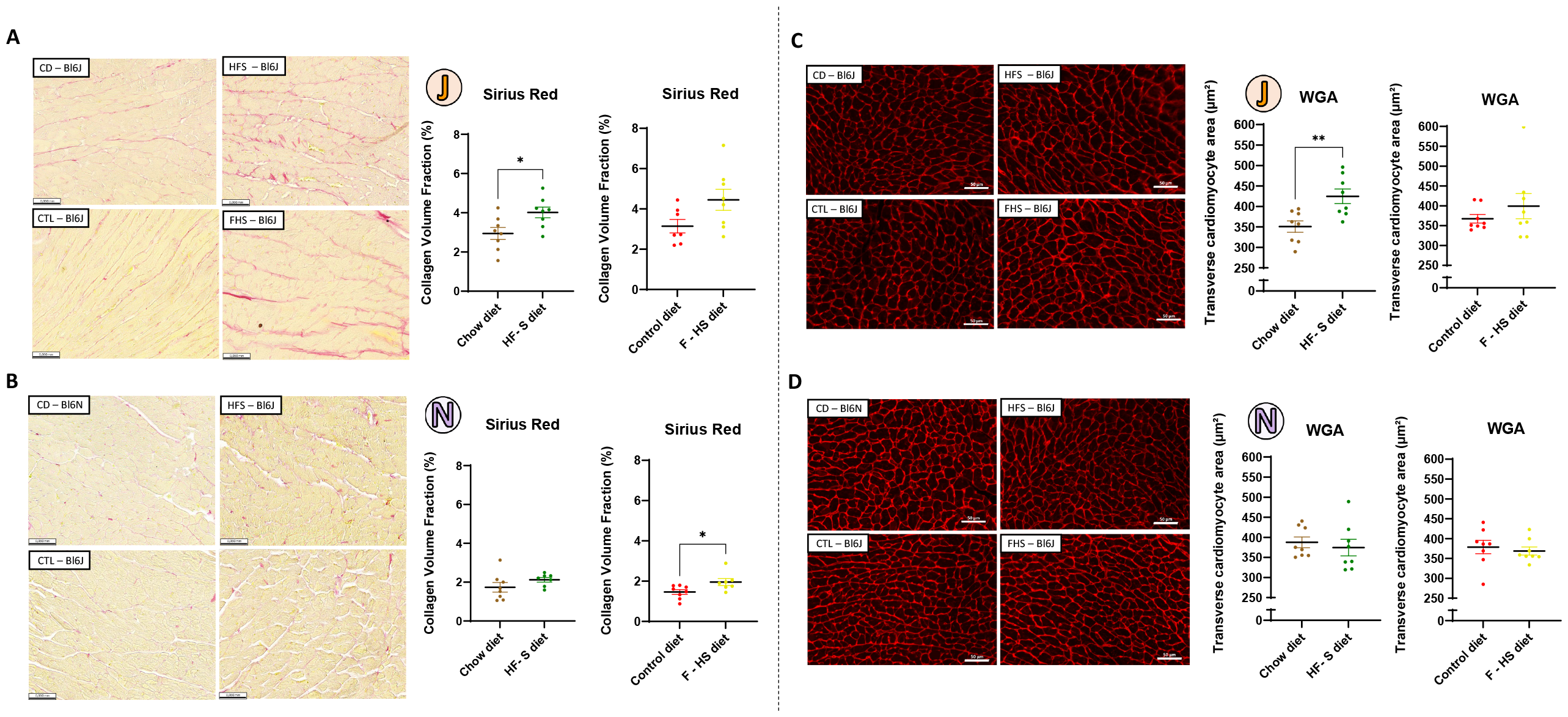
Obesogenic diet HF-S induces fibrosis and cardiac hypertrophy in C57Bl/6J but not C57Bl/6N. (A-B) Quantification of the myocardial interstitial fibrosis by Sirius Red staining (scale bar 20μm) and (C-D) by Wheat Germ Agglutinin (WGA) staining (scale bar 50μm) of the C57Bl/6J (A-C) and C57Bl/6N (B-D) mice treated with chow diet (brown), control diet (red), HF – S (green) and F – HS (yellow) (B) after 22 weeks of treatments. Data indicate the mean +/-SEM (n = 8 mice per group) and were analysed by a Student t-test

### Examination of tissue inflammation levels in adipose and cardiac tissues

In search of factors influencing this differential cardiac remodeling, we examined potential inter-strain differences in cardiac and adipose tissue inflammation, focusing on the effect of HF-S diet. Inguinal (subcutaneous) and epididymal (visceral) adipose tissues were isolated and transcriptional expression of TNFα and IL-6 (**Fig. 7 A-D**) and iNOS (**Suppl. Fig. 3A**). **Fig. 7** shows significant variability in abundance of these markers under chow (control) or HF-S diets, with no clear difference between 6J and 6N in both inguinal and epididymal adipose tissues. Multiplex analyses for inflammatory cell types in immunohistochemical sections of these adipose tissues show an increase in epididymal fat infiltration by F4/80^+^ macrophages, including CD86^+^ sub-types under HF-S in both strains (**Fig. 7 E-H; Suppl. Fig. 3C**), with more CD86^+^ in epididymal fat from 6N mice while infiltration of macrophages remained unchanged in inguinal adipose tissues in both 6N and 6J (**Suppl. Fig. 3B**). Similar analyses in LV sections of the same mice under control or HF-S showed much less inflammation in the myocardium, and no differences under HF-S or between strains (**Fig. 8A-C**).

**Figure 7.**
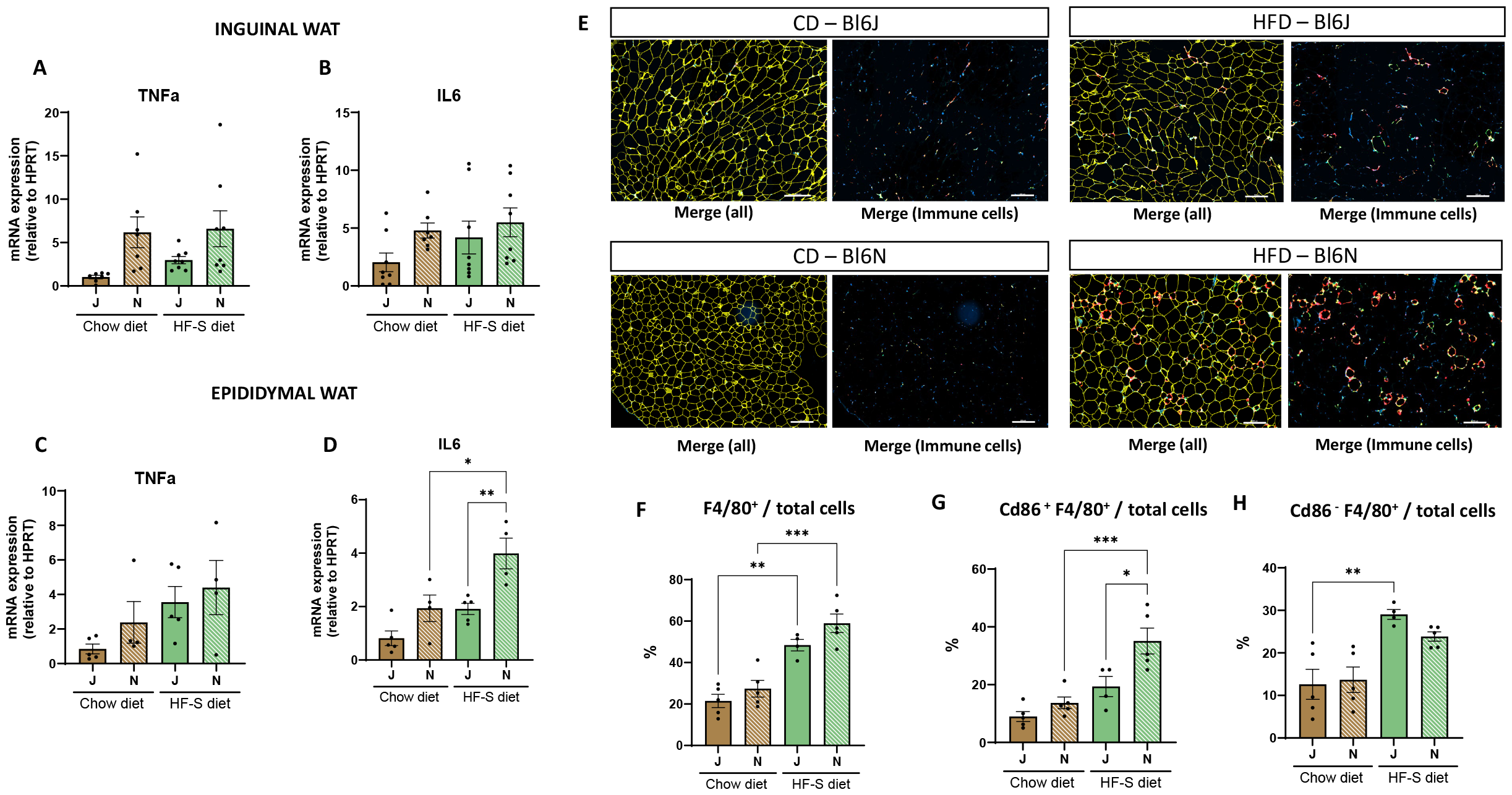
Adipose inflammatory status in C57Bl/6N and C57Bl/6J under obesogenic diets. mRNA expression of inflammatory cytokines in (A-B) inguinal or (C-D) epididymal adipose tissue of C57Bl/6J and C57Bl/6N mice treated 5 months with the chow diet (brown) or the high-fat - sucrose diet (green) (n = 6 to 8). Data indicate the mean +/-SEM and were analysed by a 2-way ANOVA. (E) Images (scale bar 200μm) and (F) proportion of macrophages (F4/80^+^) on total cells, (G) proportion of CD86^+^ macrophages (CD86^+^ and F4/80^+^) on total cells (H) proportion of CD86^-^ macrophages (CD86^-^ and F4/80^+^) on total cells by mulplex staining (CD86, F4/80) on C57Bl/6J and C57Bl/6N epididymal adipose tissue after 5 months dietary intervention with chow diet (brown) or high-fat - sucrose diet (green)(n=4 or 5 mice per group). Data indicate the mean +/-SEM and were analysed by a 2-way ANOVA

**Figure 8.**
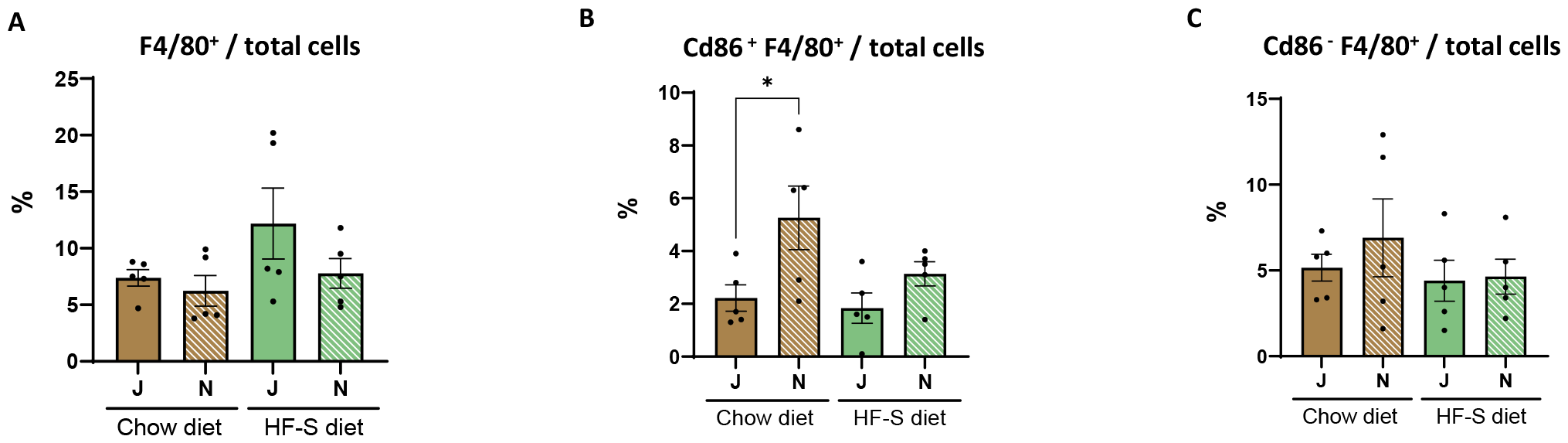
Macrophages infiltration in the le ventricle of C57Bl/6N and C57Bl/6J mice under obesogenic diets. (A) proportion of macrophages (F4/80^+^) on total cells, (B) proportion of CD86^+^ macrophages (CD86^+^ and F4/80^+^) on total cells (C) proportion of CD86^-^ macrophages (CD86^-^ and F4/80^+^) on total cells by mulplex staining (CD86, F4/80) on C57Bl/6J and C57Bl/6N le ventriclle after 5 months dietary intervention with chow diet (brown) or high-fat - sucrose diet (green). Data indicate the mean +/-SEM (n = 7 or 8 mice per group) and were analysed by a 2-way ANOVA.

## DISCUSSION

The main observations of our study are as follows; i. both HF-S and F-HS diets induce obesity and glucose intolerance in 6J and 6N mice, with similar increases in visceral and subcutaneous fat mass, adipocyte size and inflammatory infiltration; ii. 6N mice gain weight more rapidly over the first month of diet but both strains reach a similar maximal weight at M5; iii. 6J mice exhibit higher AUC for glucose after glucose load, but higher sensitivity to insulin under control diet than 6N; iv. starting at M2, both strains develop insulin resistance under obesogenic diets; v. 6N, but not 6J, mice display cardiac hypertrophic remodeling under control diets at M5, which is not further enhanced under obesogenic diets; vi. conversely, HF-S diet induces a significant increase in heart and LV hypertrophy, as well as fibrosis in 6J mice only whereas F-HS did not.

While phenotypic differences between 6N and 6J have been described under control (normal chow diet) conditions and correlations have been attempted with specific genomic variants, few studies have directly compared their response to obesogenic diets, let alone any ensuing effect on cardiac remodeling. In the present study C57Bl/6N gain weight more rapidly under obesogenic diets consistently with previous reports. However ultimately similar weight and adiposity was observed after 5 months between substrains supporting the idea that C57Bl/6J are not specifically more prone to diet-induced obesity than 6N as observed elsewhere^24^. Our observation of higher AUC in response to glucose load in 6J vs. 6N mice under control diet is in line with previous descriptions^20,22^. This difference has been attributed to a lower pancreatic beta-cell capacity to produce insulin in response to glucose as a consequence of increased oxidant stress that has been linked with genetic loss of mitochondrial NNT in 6J mice^27-29^. Accordingly, others have observed a higher constitutive oxidant stress in the 6J strain^27^. However, 6J mice retain insulin sensitivity, slightly better than 6N, under control diets as shown by the insulin tolerance tests over time. Under obesogenic diets, both strains similarly develop insulin resistance, but tend to correct their AUC for glucose by producing more insulin.

The most striking difference between the two strains is observed on cardiac remodeling, with 6J mice increasing hypertrophy and fibrosis over time under HF-S diet, whereas 6N do not. Despite loss of NNT in 6J, this is probably not related to differences in peripheral metabolism, as both strains ultimately develop a similar degree of obesity and insulin resistance under HF-S diet. Likewise, although NNT deletion has been implicated in a potentiation of the inflammatory response in immune cells^30-32^, this does not translate in any sizeable difference in inflammatory infiltrates in either adipose or heart tissues.

In cardiac cells, mitochondrial NNT was previously shown to modulate the cell response to oxidant stress generated by hemodynamic overload^27^; indeed, under higher energetic demand in stressed cells, NNT working in the “reverse mode” diverts NADPH towards NADH used in oxidative phosphorylation, thereby depleting NADPH-dependent enzymes from their substrate used for detoxification of reactive oxidant species. As a consequence, despite moderately higher systemic oxidant stress under resting conditions, 6J mice deficient in NNT are protected from excessive oxidant stress and adverse remodeling towards heart failure^28,33^. This paradigm, however, was established in response to harsh pressure overload-induced heart failure and may not be applicable to cardiometabolic disease, as in the present study.

Under careful examination, 6N mice already exhibit hypertrophic eccentric remodeling under control diets with higher heart and LV mass and larger LA area, to a level equivalent to 6J mice under HF-S diet. This probably accounts for the lack of detectable difference under HF-S in 6N mice. This progressive remodeling in absence of metabolic stress most likely results from another genetic variant specific to the 6N strain. Indeed, a previous study highlighted the presence of a mutation in the myosin light chain kinase 3 *(Mylk3)* gene in 6N mice^34^, that abolishes the expression of the protein, without compensation by other Mylk isoforms. As a consequence, 6N mice develop a mild dilated cardiomyopathy with enlarged cardiac cavities and slightly decreased LV function^34,35^. This is consistent with our observation of larger LA and increased LV mass with slightly decreased EF in 6N at M5 under control diet. Notably, this particular phenotype was shown to be independent from NNT, as it was not observed in the 6J strain, even after re-introduction of the NNT gene^34^.

Finally, the diet containing the most important proportion of lipids (HF-S; 59% Kcal from lipids, 13% Kcal from sucrose) induced cardiac remodeling over time in C57Bl/6J, whereas obesogenic diet with lower fat content (F-HS; 40% Kcal from lipids, 29% Kcal from sucrose) did not. As F-HS led in both substrains to attenuated weight gain and adiposity as well as to a delayed metabolic impairment compared to HF-S, a lower adipose tissue remodeling and reduced metabolic and inflammatory pressure exerted on the heart may well be responsible for the attenuated remodeling observed in C57Bl/6J.

## CONCLUSION

Important genetic variants in inbred strains influence the response to stress, e.g. metabolic stress under HF-S, as studied here. While NNT deletion in 6J may differetinally affect glucose tolerance, it does not influence the obese phenotype after 5 months of HF-S compared with 6N. 6J mice develop significant cardiac remodeling in response to HF-S, and seem a suitable model for cardiometabolic disease. Other genetic variants in 6N, such as in the *Mylk3* gene, produce a “constitutive” cardiomyopathy under control diet, which may obscure the effect of HF-S. This reinforces the need to carefully select the suitable mouse strain in relation to the imposed pathophysiologic stress.

## Acknowledgments

We thank D. DeMulder and R. Verdoy and N. Fabian for punctual technical support during adipocytes isolation and RNA extraction.

## Funding

This work was supported by grant *AdipoBeta3* from FRS-FNRS.

## Authors contribution

L.Y.M. and J.-L.B. conceived the original hypothesis. L.Y.M. planned the experiments. L.C., H.E. performed the core of the experiments with help from L.Y.M. and analyzed the results with input from C.D and J.-L.B. H.E. performed all echocardiographic measurements. P.M. performed the multiplex IHC under the guidance of C.B.. L.C., L.Y.M and J.-L.B wrote the manuscript. L.C. worked on illustration of the data.

## Competing interests

The authors declare that they have no competing interests.

## Data and materials availability

All data associated with this study are available in the main text or the Supplementary Materials.

## FIGURES LEGENDS

**Supplemental figure 1.**
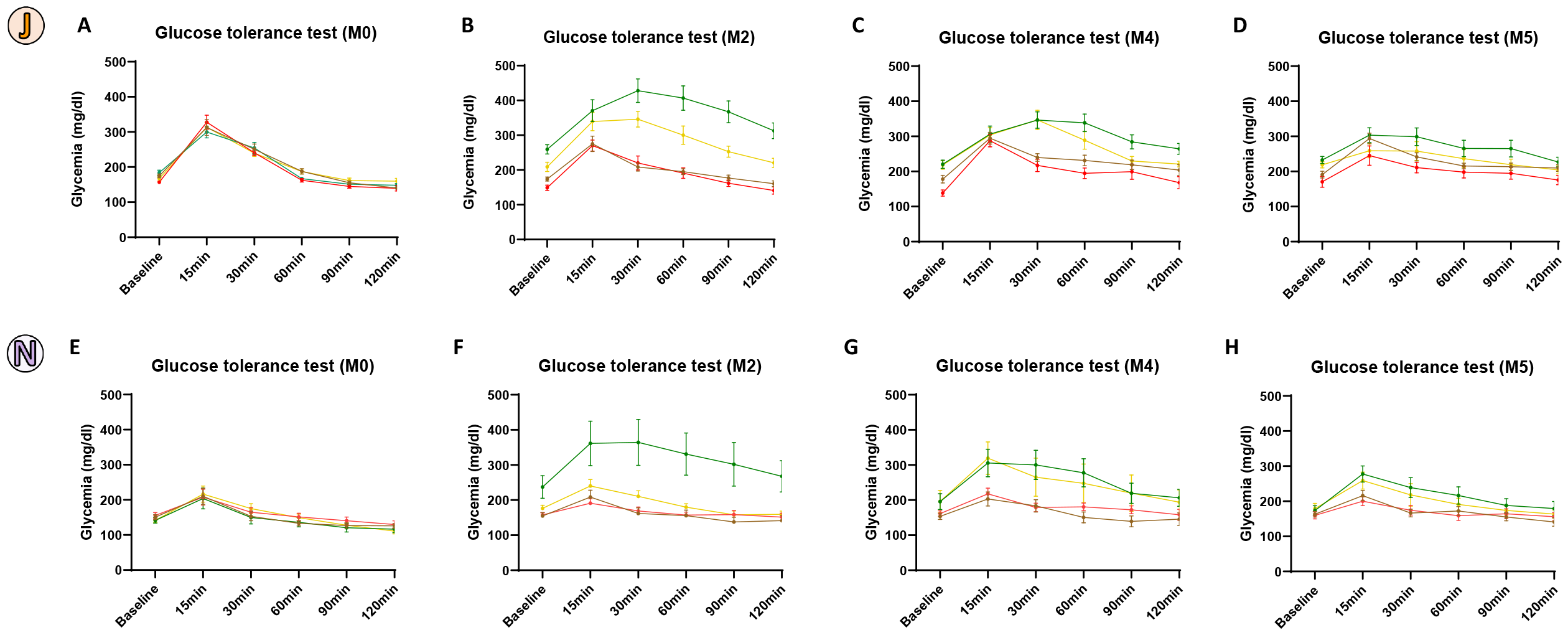
Effect of obesogenic diets on glucose tolerance in C57Bl/6 substrains. Plasma glucose during intraperitoneal glucose tolerance test (2g/kg, 6h-fasting) of the C57Bl/6J mice (A-D) and C57Bl/6N mice (E-H) treated with chow diet (brown), control diet (red), HF – S (green) and F – HS (yellow) at baseline (A-E) and after 2 (B-F), 4 (C-G) and 5 (D-H) months of treatment with the diets. Data indicate the mean +/-SEM (n = 8 mice per group) and were analysed by a 2-way repeated-measures ANOVA

**Supplemental figure 2.**
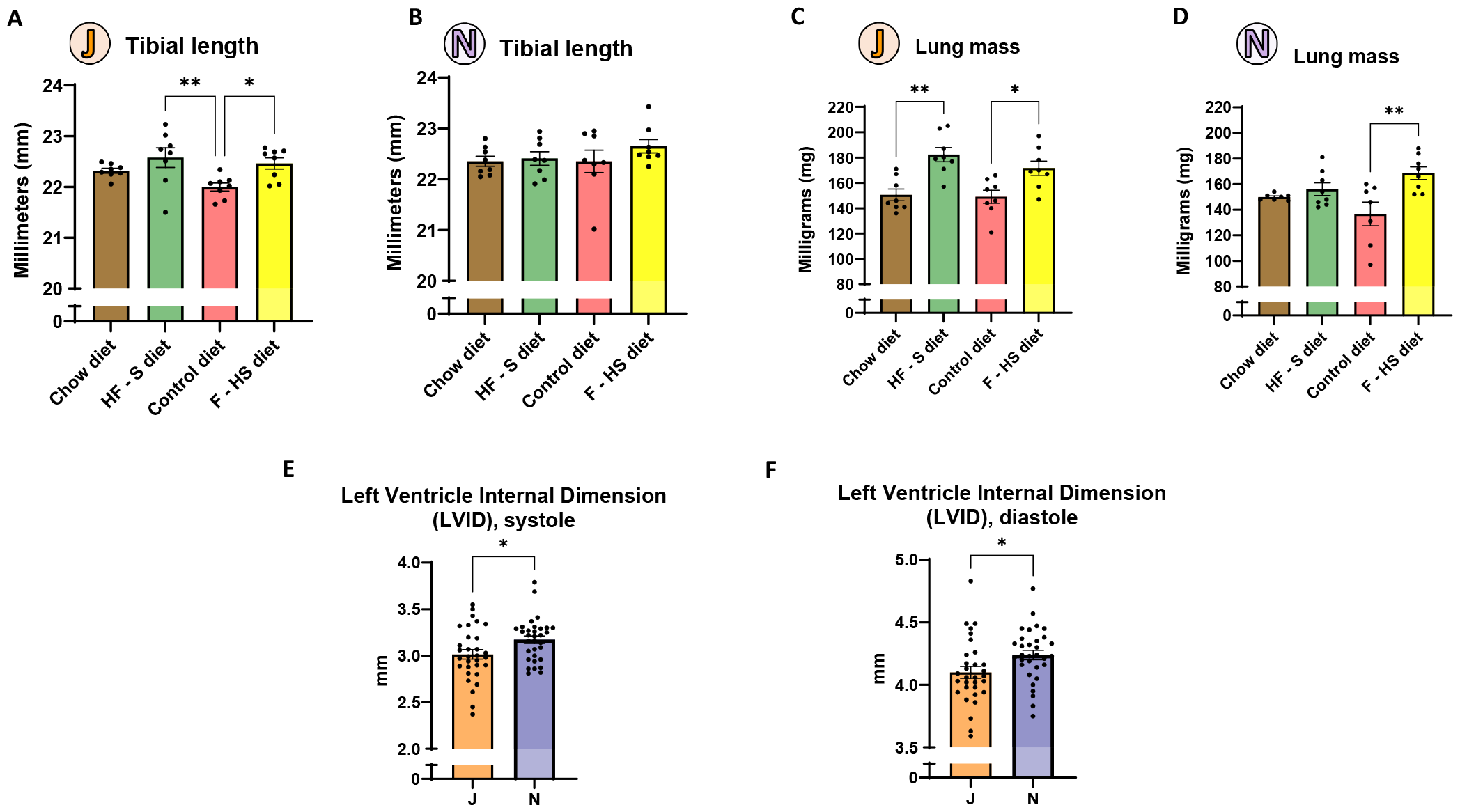
Additional morphometric and echocardiographic parameters. Tibial length of the C57Bl/6J (A) and C57Bl/6N (B) mice and lung mass of the C57Bl/6J (C) and C57Bl/6N (D) fed with chow diet (brown), control diet (red), HF – S (green) and F – HS (yellow) for 22 weeks (n = 8 mice per group).Data indicate the mean +/-SEM and were analysed by a 1-way ANOVA. Strain differences on the le ventricle internal dimension (LVID) between J and N in systole (E) and in diastole (F). Data indicate the mean +/-SEM (n = 32 mice per group) and were analysed by a Student t-test.

**Supplemental figure 3.**
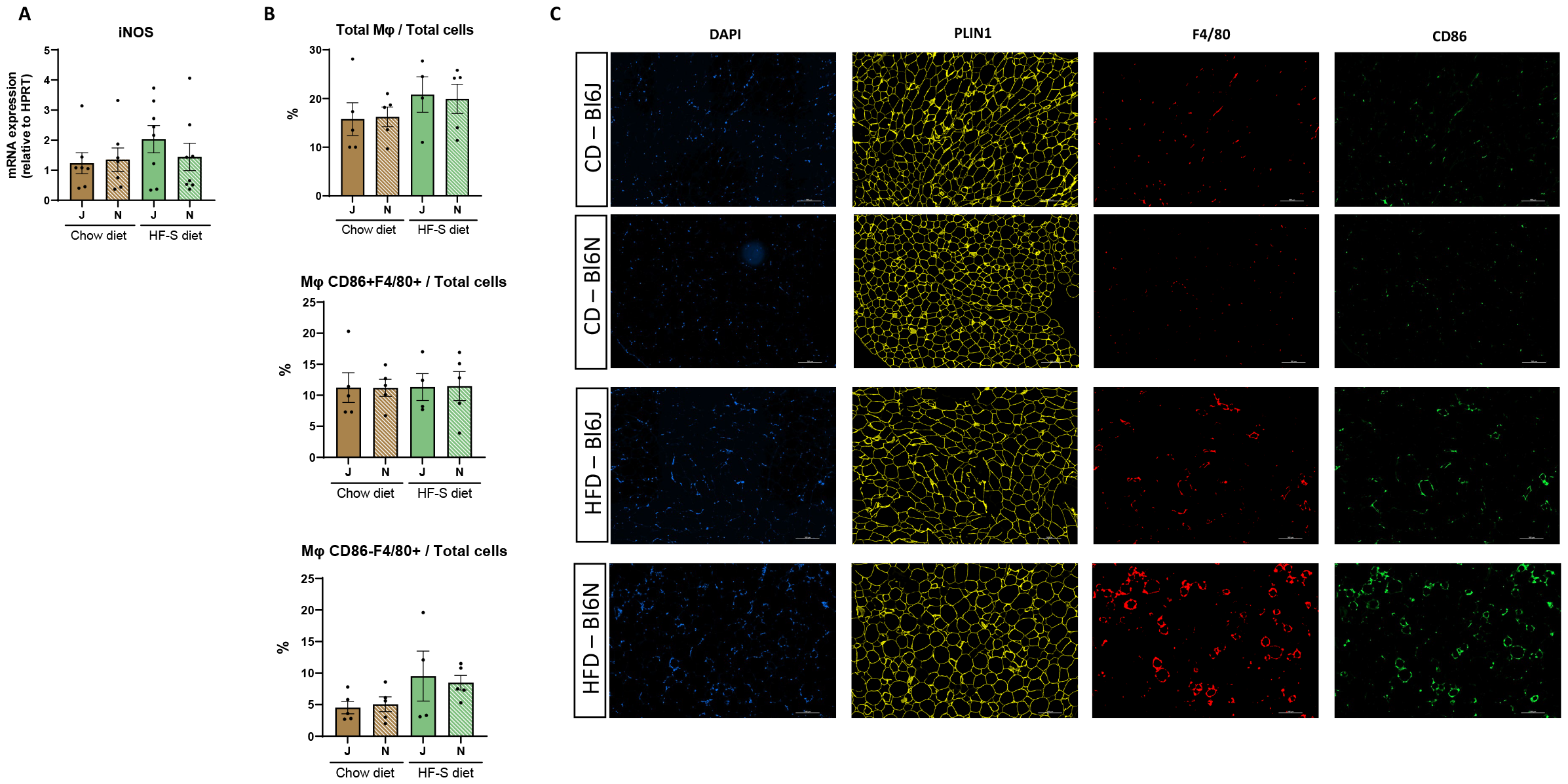
Inflammation in adipose tissue under obesogenic diets. (A) mRNA expression of inflammation marker iNOS in inguinal adipose tissue of C57Bl/6J and C57Bl/6N mice treated 5 months with the chow diet (brown) or the high-fat - sucrose diet (green) (n = 6 to 8). (B) proportion of macrophages (F4/80^+^) on total cells, proportion of CD86^+^ macrophages (CD86^+^ and F4/80^+^) on total cells; proportion of CD86^-^ macrophages (CD86^-^ and F4/80^+^) on total cells by multiplex staining (CD86, F4/80) on C57Bl/6J and C57Bl/6N inguinal adipose tissues after 5 months dietary intervention with chow diet (brown) or high-fat - sucrose diet (green)(n=4 or 5 mice per group). Data indicate the mean +/-SEM and were analysed by a 2-way ANOVA. (C) representative images of of separate staining in epididymal adipose tissue for CD86, F4/80, perilipin-1 and Dapi as showed combined in figure 7.

## Notes

### Competing Interest Statement

The authors have declared no competing interest.

## REFERENCES

1. Saklayen, M.G. (2018). The Global Epidemic of the Metabolic Syndrome. Curr Hypertens Rep 20, 12. 10.1007/s11906-018-0812-z.

2. Hossain, P., Kawar, B., and El Nahas, M. (2007). Obesity and diabetes in the developing world--a growing challenge. The New England journal of medicine 356, 213–215. 10.1056/NEJMp068177.

3. Grundy, S.M., Cleeman, J.I., Daniels, S.R., Donato, K.A., Eckel, R.H., Franklin, B.A., Gordon, D.J., Krauss, R.M., Savage, P.J., Smith, S.C., Jr., et al. (2005). Diagnosis and management of the metabolic syndrome: an American Heart Association/National Heart, Lung, and Blood Institute Scientific Statement. Circulation 112, 2735–2752. 10.1161/CIRCULATIONAHA.105.169404.

4. Chobanian, A.V., Bakris, G.L., Black, H.R., Cushman, W.C., Green, L.A., Izzo, J.L., Jr., Jones, D.W., Materson, B.J., Oparil, S., Wright, J.T., Jr., et al. (2003). The Seventh Report of the Joint National Committee on Prevention, Detection, Evaluation, and Treatment of High Blood Pressure: the JNC 7 report. Jama 289, 2560–2572. 10.1001/jama.289.19.2560.

5. Authors/Task Force, M., Ryden, L., Grant, P.J., Anker, S.D., Berne, C., Cosentino, F., Danchin, N., Deaton, C., Escaned, J., Hammes, H.P., et al. (2013). ESC Guidelines on diabetes, pre-diabetes, and cardiovascular diseases developed in collaboration with the EASD: the Task Force on diabetes, pre-diabetes, and cardiovascular diseases of the European Society of Cardiology (ESC) and developed in collaboration with the European Association for the Study of Diabetes (EASD). European heart journal 34, 3035–3087. 10.1093/eurheartj/eht108.

6. Ng, A.C.T., Delgado, V., Borlaug, B.A., and Bax, J.J. (2021). Diabesity: the combined burden of obesity and diabetes on heart disease and the role of imaging. Nat Rev Cardiol 18, 291–304. 10.1038/s41569-020-00465-5.

7. Kitzman, D.W., and Lam, C.S.P. (2017). Obese Heart Failure With Preserved Ejection Fraction Phenotype: From Pariah to Central Player. Circulation 136, 20–23. 10.1161/CIRCULATIONAHA.117.028365.

8. Tromp, J., Claggett, B.L., Liu, J., Jackson, A.M., Jhund, P.S., Kober, L., Widimsky, J., Boytsov, S.A., Chopra, V.K., Anand, I.S., et al. (2021). Global Differences in Heart Failure With Preserved Ejection Fraction: The PARAGON-HF Trial. Circulation. Heart failure 14, e007901. 10.1161/CIRCHEARTFAILURE.120.007901.

9. Schiattarella, G.G., Alcaide, P., Condorelli, G., Gillette, T.G., Heymans, S., Jones, E.A.V., Kallikourdis, M., Lichtman, A., Marelli-Berg, F., Shah, S., et al. (2022). Immunometabolic Mechanisms of Heart Failure with Preserved Ejection Fraction. Nat Cardiovasc Res 1, 211–222. 10.1038/s44161-022-00032-w.

10. Obokata, M., Reddy, Y.N.V., Pislaru, S.V., Melenovsky, V., and Borlaug, B.A. (2017). Evidence Supporting the Existence of a Distinct Obese Phenotype of Heart Failure With Preserved Ejection Fraction. Circulation 136, 6–19. 10.1161/CIRCULATIONAHA.116.026807.

11. Mentz, R.J., Kelly, J.P., von Lueder, T.G., Voors, A.A., Lam, C.S., Cowie, M.R., Kjeldsen, K., Jankowska, E.A., Atar, D., Butler, J., et al. (2014). Noncardiac comorbidities in heart failure with reduced versus preserved ejection fraction. Journal of the American College of Cardiology 64, 2281–2293. 10.1016/j.jacc.2014.08.036.

12. Keane, T.M., Goodstadt, L., Danecek, P., White, M.A., Wong, K., Yalcin, B., Heger, A., Agam, A., Slater, G., Goodson, M., et al. (2011). Mouse genomic variation and its effect on phenotypes and gene regulation. Nature 477, 289–294. 10.1038/nature10413.

13. Montgomery, M.K., Hallahan, N.L., Brown, S.H., Liu, M., Mitchell, T.W., Cooney, G.J., and Turner, N. (2013). Mouse strain-dependent variation in obesity and glucose homeostasis in response to high-fat feeding. Diabetologia 56, 1129–1139. 10.1007/s00125-013-2846-8.

14. Fearnside, J.F., Dumas, M.E., Rothwell, A.R., Wilder, S.P., Cloarec, O., Toye, A., Blancher, C., Holmes, E., Tatoud, R., Barton, R.H., et al. (2008). Phylometabonomic patterns of adaptation to high fat diet feeding in inbred mice. PloS one 3, e1668. 10.1371/journal.pone.0001668.

15. Hull, R.L., Willard, J.R., Struck, M.D., Barrow, B.M., Brar, G.S., Andrikopoulos, S., and Zraika, S. (2017). High fat feeding unmasks variable insulin responses in male C57BL/6 mouse substrains. The Journal of endocrinology 233, 53–64. 10.1530/JOE-16-0377.

16. Russell, E.S. (1978). Genetic origins and some research uses of c57bl/6, dba/2, and b6d2f1 mice. In Development of the rodent as a model system of aging, A.R. Gibson DC, Finch C, ed. pp. 37–44.

17. Fontaine, D.A., and Davis, D.B. (2016). Attention to Background Strain Is Essential for Metabolic Research: C57BL/6 and the International Knockout Mouse Consortium. Diabetes 65, 25–33. 10.2337/db15-0982.

18. Festing, M.F. (1996). Origins and characteristics of inbred strains of mice. In Genetic variants and strains of the laboratory mouse, R.S. Lyon MF, Brown SDM, ed. pp. 1537–1576.

19. Simon, M.M., Greenaway, S., White, J.K., Fuchs, H., Gailus-Durner, V., Wells, S., Sorg, T., Wong, K., Bedu, E., Cartwright, E.J., et al. (2013). A comparative phenotypic and genomic analysis of C57BL/6J and C57BL/6N mouse strains. Genome Biol 14, R82. 10.1186/gb-2013-14-7-r82.

20. Kaku, K., Fiedorek, F.T., Jr., Province, M., and Permutt, M.A. (1988). Genetic analysis of glucose tolerance in inbred mouse strains. Evidence for polygenic control. Diabetes 37, 707–713. 10.2337/diab.37.6.707.

21. Lee, S.K., Opara, E.C., Surwit, R.S., Feinglos, M.N., and Akwari, O.E. (1995). Defective glucose-stimulated insulin release from perifused islets of C57BL/6J mice. Pancreas 11, 206–211. 10.1097/00006676-199508000-00016.

22. Toye, A.A., Lippiat, J.D., Proks, P., Shimomura, K., Bentley, L., Hugill, A., Mijat, V., Goldsworthy, M., Moir, L., Haynes, A., et al. (2005). A genetic and physiological study of impaired glucose homeostasis control in C57BL/6J mice. Diabetologia 48, 675–686. 10.1007/s00125-005-1680-z.

23. Nicholson, A., Reifsnyder, P.C., Malcolm, R.D., Lucas, C.A., MacGregor, G.R., Zhang, W., and Leiter, E.H. (2010). Diet-induced obesity in two C57BL/6 substrains with intact or mutant nicotinamide nucleotide transhydrogenase (Nnt) gene. Obesity (Silver Spring) 18, 1902–1905. 10.1038/oby.2009.477.

24. Fisher-Wellman, K.H., Ryan, T.E., Smith, C.D., Gilliam, L.A., Lin, C.T., Reese, L.R., Torres, M.J., and Neufer, P.D. (2016). A Direct Comparison of Metabolic Responses to High-Fat Diet in C57BL/6J and C57BL/6NJ Mice. Diabetes 65, 3249–3261. 10.2337/db16-0291.

25. Khisti, R.T., Wolstenholme, J., Shelton, K.L., and Miles, M.F. (2006). Characterization of the ethanol-deprivation effect in substrains of C57BL/6 mice. Alcohol 40, 119–126. 10.1016/j.alcohol.2006.12.003.

26. Ulmasov, B., Oshima, K., Rodriguez, M.G., Cox, R.D., and Neuschwander-Tetri, B.A. (2013). Differences in the degree of cerulein-induced chronic pancreatitis in C57BL/6 mouse substrains lead to new insights in identification of potential risk factors in the development of chronic pancreatitis. Am J Pathol 183, 692–708. 10.1016/j.ajpath.2013.05.020.

27. Ronchi, J.A., Figueira, T.R., Ravagnani, F.G., Oliveira, H.C., Vercesi, A.E., and Castilho, R.F. (2013). A spontaneous mutation in the nicotinamide nucleotide transhydrogenase gene of C57BL/6J mice results in mitochondrial redox abnormalities. Free Radic Biol Med 63, 446–456. 10.1016/j.freeradbiomed.2013.05.049.

28. Nickel, A.G., von Hardenberg, A., Hohl, M., Loffler, J.R., Kohlhaas, M., Becker, J., Reil, J.C., Kazakov, A., Bonnekoh, J., Stadelmaier, M., et al. (2015). Reversal of Mitochondrial Transhydrogenase Causes Oxidative Stress in Heart Failure. Cell metabolism 22, 472–484. 10.1016/j.cmet.2015.07.008.

29. Freeman, H.C., Hugill, A., Dear, N.T., Ashcroft, F.M., and Cox, R.D. (2006). Deletion of nicotinamide nucleotide transhydrogenase: a new quantitive trait locus accounting for glucose intolerance in C57BL/6J mice. Diabetes 55, 2153–2156. 10.2337/db06-0358.

30. Salerno, A.G., Rentz, T., Dorighello, G.G., Marques, A.C., Lorza-Gil, E., Wanschel, A., de Moraes, A., Vercesi, A.E., and Oliveira, H.C.F. (2019). Lack of mitochondrial NADP(H)-transhydrogenase expression in macrophages exacerbates atherosclerosis in hypercholesterolemic mice. The Biochemical journal 476, 3769–3789. 10.1042/BCJ20190543.

31. Ripoll, V.M., Meadows, N.A., Bangert, M., Lee, A.W., Kadioglu, A., and Cox, R.D. (2012). Nicotinamide nucleotide transhydrogenase (NNT) acts as a novel modulator of macrophage inflammatory responses. FASEB J 26, 3550–3562. 10.1096/.11-199935.

32. Regan, T., Conway, R., and Bharath, L.P. (2022). Regulation of immune cell function by nicotinamide nucleotide transhydrogenase. American journal of physiology. Cell physiology 322, C666–C673. 10.1152/ajpcell.00607.2020.

33. Garcia-Menendez, L., Karamanlidis, G., Kolwicz, S., and Tian, R. (2013). Substrain specific response to cardiac pressure overload in C57BL/6 mice. American journal of physiology. Heart and circulatory physiology 305, H397–402. 10.1152/ajpheart.00088.2013.

34. Williams, J.L., Paudyal, A., Awad, S., Nicholson, J., Grzesik, D., Botta, J., Meimaridou, E., Maharaj, A.V., Stewart, M., Tinker, A., et al. (2020). Mylk3 null C57BL/6N mice develop cardiomyopathy, whereas Nnt null C57BL/6J mice do not. Life Sci Alliance 3. 10.26508/lsa.201900593.

35. Moreth, K., Fischer, R., Fuchs, H., Gailus-Durner, V., Wurst, W., Katus, H.A., Bekeredjian, R., and Hrabe de Angelis, M. (2014). High-throughput phenotypic assessment of cardiac physiology in four commonly used inbred mouse strains. J Comp Physiol B 184, 763–775. 10.1007/s00360-014-0830-3.

